# Tracing the Transitions from Pluripotency to Germ Cell Fate with CRISPR Screening

**DOI:** 10.1101/269811

**Authors:** Jamie A. Hackett, Yun Huang, Ufuk Günesdogan, Kristjan Holm-Gretarsson, Toshihiro Kobayashi, M. Azim Surani

## Abstract

Early mammalian development entails a series of cell fate transitions that includes transit through naïve pluripotency to post-implantation epiblast. This subsequently gives rise to primordial germ cells (PGC), the founding population of the germline lineage. To investigate the gene regulatory networks that control these critical cell fate decisions, we developed a compound-reporter system to track cellular identity in a model of PGC specification (PGC-like cells; PGCLC), and coupled it with unbiased genome-wide CRISPR screening. This enabled identification of key genes both for exit from pluripotency and for acquisition of PGC fate, with further characterisation revealing a central role for the transcription factors *Nr5a2* and *Zfp296* in germline ontogeny. Abrogation of these genes results in significantly impaired PGCLC development due to widespread activation (*Nr5a2*^−/−^) or inhibition (*Zfp296*^−/−^) of WNT pathway components. This leads to aberrant upregulation of the somatic programme or failure to appropriately activate germline genes in PGCLC, respectively, and consequently loss of germ cell identity. Overall our study places *Zfp296* and *Nr5a2* as key components of an expanded PGC gene regulatory network, and outlines a transferable strategy for identifying critical regulators of complex cell fate transitions.

## INTRODUCTION

The germ cell lineage generates the totipotent state and transmits heritable genetic and epigenetic information to the next generation. Robust specification of primordial germ cells (PGC), the precursors of sperm and eggs, is therefore a critical developmental event to ensure the propagation of a species. Approximately thirty specified PGCs are detected in mouse embryos at embryonic day (E) 7.25, which arise from ‘competent’ epiblast precursor cells in response to BMP and WNT signalling^1,2^. The specification of PGCs is accompanied by induction of key germ-cell genes, repression of the nascent somatic-mesodermal programme, and widespread epigenetic remodelling, including global DNA demethylation^3,4^.

PGC specification follows WNT-dependent induction of the primitive streak/mesodermal gene *Brachyury (T)*, which in mice promotes activation of key PGC specifiers; *Blimp1, Prdm14, Cbfa2t2*, and *Ap2*γ^5–7^. These transcription factors form a self-reinforcing network that feeds back to supress other WNT/BMP-induced mesodermal genes (including *T*), thereby repressing the ongoing somatic programme in early PGCs^8^. *Blimp1* and *Prdm14* additionally activate germline-specific genes and initiate epigenome resetting, with mutation of either gene resulting in loss of PGCs by E12.5^9,10^. The broader gene regulatory network that controls PGC ontogeny has however been relatively uncharted, due to the absence of unbiased functional approaches and the challenges of analysing the limited number of nascent PGCs in embryos.

The recent development of an *in vitro* derived model of PGCs, termed PGC-like cells (PGCLC), now facilitates molecular studies of the specification and early differentiation events of germ cell fate^11^. PGCLC are derived by inducing naïve embryonic stem cells (ESC) that are equivalent to the inner cell mass (ICM) of blastocysts (E3.5-E4.0), towards competent epiblast-like cells (EpiLC), which closely resemble pre-gastrulation mouse epiblast (E5.5-E6.5)^12^. EpiLC can in turn be induced to undergo specification as PGCLC in response to BMP and WNT. Specified PGCLC are equivalent to migratory PGCs *in vivo* (E8.5-E10.5)^11^, and have the potential to develop to mature functional gametes^13,14^. This model is therefore appropriate to investigate the inherent regulatory mechanisms of nascent germ cells, and the preceding developmental transitions from ICM, through post-implantation development and PGC specification.

The advent of genome-wide CRISPR screening has enabled unbiased interrogation of recessive gene function in a wide spectrum of biological contexts^15^. We reasoned that by designing appropriate reporters, CRISPR screening could be adapted to identify genes involved in controlling successive cell fate decisions during lineage-specific differentiation. Specifically, by employing the PGCLC model we embarked on identification of genes that are important for (i) maintenance of naïve pluripotency, (ii) transition to the germline competent state (EpiLC) and, (iii) specification of the PGC lineage. We further investigate novel candidates to reveal their key role and mechanistic function during nascent germ cell development. From a broader perspective, we demonstrate that unbiased CRISPR screening can be adapted to probe the genetic basis of complex multi-step developmental processes, using the germline lineage as a paradigm.

## RESULTS

### A compound-reporter for developmental transitions toward PGC fate

We set out to design a reporter system that can distinguish between successive cell identities during the developmental transitions from naïve pluripotency to specified primordial germ cell (PGC) fate. Single-cell RNA-seq data revealed that *Stella* (also known as *Dppa3*) is expressed in the naïve pluripotent ICM but is rapidly downregulated in postimplanation epiblast, with re-activation occurring specifically in nascent PGCs (from E7.5)^16^. In contrast *Esg1* (also known as *Dppa5*) is also expressed in the ICM, but maintains high expression during postimplanation development, and subsequently becomes strongly downregulated in early PGCs; with low germline expression reacquired later (>E9.5) (Fig 1A). To exploit the mutually exclusive expression of these genes, we generated an ESC line with compound *<u>S</u>tella*-<u>G</u>FP and *<u>E</u>sg1*-td<u>T</u>omato (SGET) reporters. In this ‘traffic-light’ system naïve pluripotent cells are double-positive (yellow), postimplantation epiblast cells are *Esg1*-positive (red), and nascent PGCs are *Stella*-positive (green). We monitored SGET expression during mouse development by tetraploid complementation and chimera formation, and observed strong double-activation in naïve pluripotent epiblast at E4.5 (Fig 1B). This resolved to single *Esg1-tdTomato* activity in E6.0 epiblast, with both reporters subsequently silenced in somatic tissues by E7.0. Importantly *Stella*-GFP is specifically reactivated in PGCs by E8.5, until E12.5, with *Esg1*-tdTomato additionally showing weak expression in later PGC stages (Fig 1B). Thus, SGET faithfully recapitulates expression of the endogenous genes during the transitions towards PGC fate.

**Figure 1.**
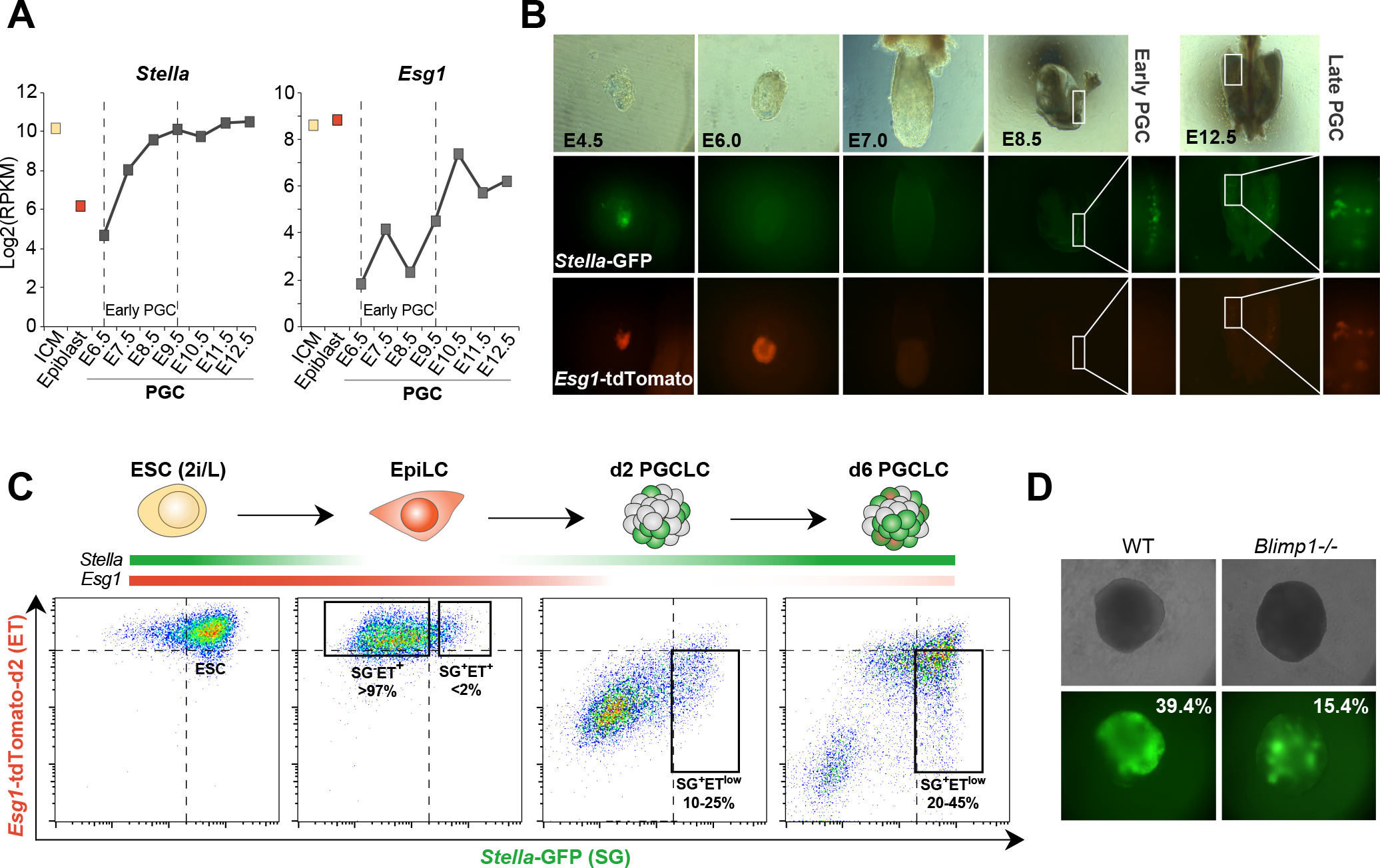
The SGET reporter for tracking cell fate transitions toward PGCs. **(A)** Single-cell RNA-seq analysis showing reciprocal gene expression patterns of endogenous *Stella* and *Esg1* during early development and in primordial germ cells (PGC). **(B)** Chimera and tetraploid complementation assays confirm faithful expression of the *Stella*-GFP (green) and Esg7-tdTomato (red; SGET) reporter during early development and in PGCs. **(C)** Representative FACS analysis of SGET activation during *in vitro* cell fate transitions of ESC into EpiLC and subsequently PGCLC. **(D)** Representative images of impaired PGCLC induction from *Blimp1*^−/−^ SGET ESC. Numbers indicate percentage of SG^+^ET^low^ at day 5 as determined by FACS.

Next, we examined SGET activity during *in vitro* specification of PGCLC, which initiates from naïve ESC and transitions through ‘competent’ epiblast-like cells (EpiLC)^11^. SGET ESC were largely double-positive (SG^+^ET^+^), but resolved to *Esg1*-only expression (SG^−^ET^+^) in >97% of cells upon induction into EpiLC. In contrast, nascent PGCLC (day 2) reactivated *Stella* whilst concomitantly repressing *Esg1* (SG^+^ET^−^), with later stage PGCLC (day 6) exhibiting low *Esg1* expression (SG^+^ET^low^) (Fig 1C). PGCLC carrying SGET therefore recapitulate the *in vivo* dynamics. To functionally validate the SGET reporter system, we generated ESC with a mutation in the key PGC specifier *Blimp1* (Fig S1A), which resulted in a significant reduction (up to 2.8-fold) in the efficiency of PGCLC generation (Fig 1D & S1B). The remaining *Blimp1*^−/−^ PGCLCs exhibited aberrant gene expression (Fig S1C), consistent with *in vivo* studies where a fraction of nascent ‘PGCs’ remain but with an altered transcriptome^9^. The SGET reporter system is thus ideal for monitoring successive changes in cell identity, from naïve ESCs to specified PGCLC.

### CRISPR screen identifies key genes for naïve ESC

We introduced a single-copy of *Cas9* into SGET ESC (Fig S1D), and subsequently infected this line with an integrating lentiviral library of exon-targeting guide RNAs (gRNA)^17^, in independent biological replicates. Statistical analysis of gRNA frequency in the ESC population, relative to the initial frequency, reveals essential genes because their cognate gRNAs become depleted concomitant with cell loss. We reasoned that by flow sorting SGET cells that successfully acquire each sequential fate during PGCLC specification, we could identify essential genes for each transition by comparing gRNA frequency to the preceding cell population (Fig S1E). Accordingly, this approach filters out genes with a role in prior fate transitions, and reveals the critical regulators for each stage of multi-step differentiation events, based on functional requirement.

Following introduction of the CRISPR library, we identified 627 genes that lead to a loss of naïve ESC maintained in 2i/LIF when knocked-out (*p*<0.01 in both replicates) (Fig 2A). Gene ontology (GO) terms for these genes were highly enriched for fundamental biological processes such as ribosome biogenesis (*p*=2.8 × 10^−27^; e.g. *Rps5*) and protein translation (*p*= 4.2 × 10^−30^; e.g. *Eif6*), supporting the efficacy of the library (Fig S2A). Notably ESC carrying mutations for the core pluripotency factors *Oct4* and *Sox2* were also highly depleted, consistent with their essential role in propagating pluripotent ESC^18^ (Fig 2A, B; S2C). The majority of naïve pluripotency genes were not depleted however, which is in line with the capacity of 2i/LIF media to buffer against a single genetic perturbation to the naïve pluripotency network (Fig 2B)^19^. Importantly, this enables the role of such naïve pluripotency genes to be examined in PGC specification, since knockout cells remain in the population. Exceptions however are *Nanog* and *Tfcp2l1*, which are likely depleted owing to a proliferation disadvantage upon knockout in ESC. In contrast to the many depleted genes, we observed that only twenty-one genes become enriched in the population upon knockout, primarily corresponding to tumour-suppressors (Fig S2B & C). Amongst these *Trp53* (*p53*) is the top hit, suggesting that *p53-*mutant mouse ESC acquire a major selective advantage, similarly to recent reports in human ESC^20^.

**Figure 2.**
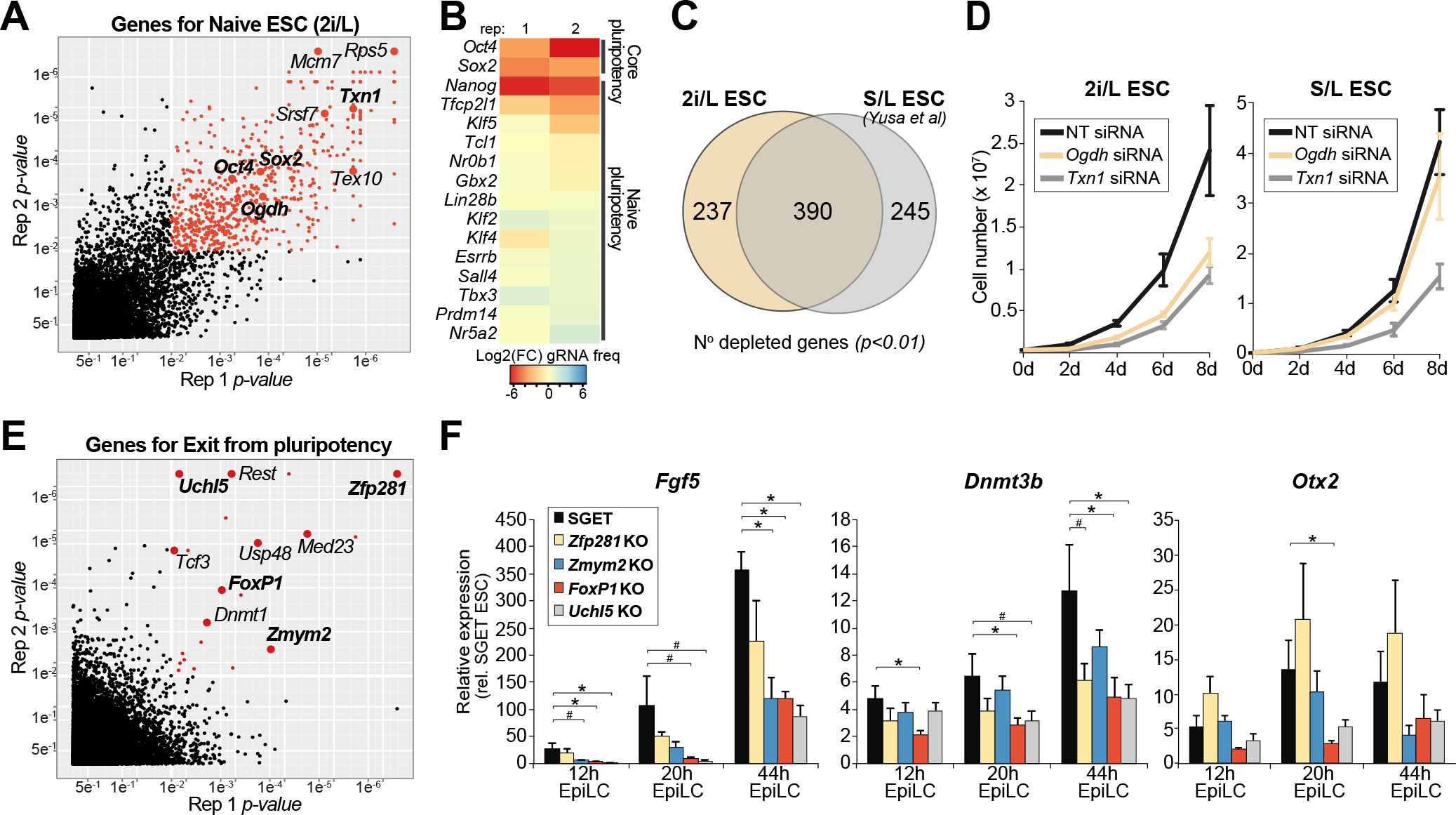
Identification of important genes for ESC and EpiLC induction. **(A)** Scatter plot showing essential genes for ESC propagation in 2i/L (red datapoints) as determined by a CRISPR screen. **(B)** Heamap showing mean fold change (FC) in normalised frequency of gRNAs targeting core-and naïve- pluripotency genes in ESC relative to frequency in starting gRNA library. Reduced frequency indicates a functionally important gene for ESC in 2i/L. **(C)** Venn diagram intersecting significant depleted genes in 2i/L ESC with serum/LIF (S/L) maintained ESC. **(D)** Proliferation of ESC following siRNA knockdown of *Ogdh* or *Txn1*in ESC maintained in 2i/L or S/L culture conditions, respectively. **(E)** Scatter plot showing significantly enriched gene knockouts in SG^+^ET^+^ EpiLC that have failed to exit naïve pluripotency. **(F)** Gene expression showing impaired activation of key epiblast markers in EpiLC carrying knockouts for candidate exit from pluripotency regulators. Significance was determined using a one-tailed T-test; * *p*<*0.05*; ^#^ *p*<*0.1*. Error bars show s.e.m of triplicate independent experiments.

We next compared the genes essential for naïve ESC (in 2i/LIF) with those essential for ESC in conventional serum/LIF culture, which represents an alternative pluripotent state^18^. Using datasets from the same CRISPR library^17^, we observed good overlap (62%) between ESC conditions, but noted 38% of genes appear to be critical under only specific pluripotent culture parameters (Fig 2C). Among the 2i/LIF specific genes were *Ogdh* and *Dlst*; two enzymes that mediate α-ketoglutarate metabolism which feeds into the TCA cycle. We used siRNAs to test acute loss of *Ogdh* and observed a significant reduction of proliferation and/or increased cell death in 2i/LIF ESC but not serum/LIF ESC (Fig 2D). We also tested *Txn1*, which was scored as essential in both pluripotent ESC states, and found an equivalent cell reduction in 2i/LIF and serum/LIF (Fig 2D). These data suggest ESC in distinct pluripotent states rely, in part, on distinct genetic networks, with ESC in 2i/L uniquely reliant on *Ogdh* metabolism of α-ketoglutarate for example. More generally, establishing this mutant SGET ESC population enables screening for functional regulators of EpiLCs, and subsequently PGCLCs, without confounding general survival genes.

### Acquisition of post-implantation epiblast fate

We next used fluorescence-activated flow sorting (FACS) to isolate EpiLC that successfully transited to the SG^−^ET^+^ epiblast state, as well as cells that maintained *Stella* expression (SG^+^ET^+^) (<2%), indicative of failure to exit naïve pluripotency. Analysis of this SG^+^ET^+^ EpiLC population revealed enrichment for 21 candidate genes with potential intrinsic roles in dissolution of naïve pluripotency (*p*<*0.01*), including the two prototypical regulators *Tcf3* and *Zfp281* (Fig 2E & S3A)^21,22^. The candidates additionally included the epigenetic regulator *Dnmt1* and also *Rest*, which are known to be important during early differentiation^23,24^.

To test the role of novel candidates in driving exit from pluripotency we generated ESC knockouts for *Zmym2 (*also known as *Zfp198), FoxP1*, *Uchl5*, and *Zfp281* as a positive control, using CRISPR targeting. Consistent with their failure to repress *Stella*-GFP during EpiLC formation, mutant EpiLC were also impaired in activation of key epiblast markers *Fgf5*, *Dnmt3b* and *Otx2*, whilst their proliferation appeared unaffected (Fig 2F & S3B-C). We observed both a delay in activation of epiblast genes and a reduction in their absolute levels in knockout lines, despite high levels of powerful differentiation-inducing FGF and ACTIVIN in the culture medium. Notably, the *Zmym2*-, *FoxP1*-, *Uchl5*- mutant lines exhibited expression defects comparable with the *bona fide* exit from pluripotency regulator *Zfp281*, supporting an important role for them in the timely acquisition of post-implantation cell identity. Consistently all three candidates are transcriptionally upregulated during EpiLC induction (Fig S3D).

### Candidate genes for specification of PGC fate

We next focused on genes involved in mouse germline specification by inducing PGCLC from competent SG^−^ET^+^ EpiLC. In order to obtain sufficient numbers of PGCLCs for coverage of the gRNA library, we optimised and scaled-up the induction (Fig S4A). Specified SG^+^ET^low^ PGCLC were FACS purified at d6 to allow time for depletion of cells carrying mutations in critical germline genes. Because PGCLC numbers remained a limiting factor however, we relaxed the candidate threshold to *p*<*0.05*, whilst retaining a *p*<*0.01* in at least one replicate, to account for increased noise (Fig S4B). This resulted in identification of 23 candidate genes involved in specification and/or development of nascent PGC. The frequency of gRNAs targeting these genes was highly depleted in PGCLC relative to preceding EpiLC (Fig 3A), whilst control gene families were not depleted (Fig S4C). Moreover, genes with an established critical role in PGC specification, such as *Blimp1, Cbfat2t2* and *Prdm14*, also showed marked reduction in gRNA frequency in PGCLC, supporting the efficacy of the screen (Fig 3A). These genes narrowly failed to meet significance across all replicates however; *Cbfa2t2* scored p-values of *p*=*0.00006* and *p*=*0.22* for example. We also noted that several pluripotency genes (*Nr5a2*, *Esrrb* and *Sall4*) were depleted specifically from PGCLC (Fig S4C), suggesting several genes typically linked with pluripotency have a potentially important function in the germline.

**Figure 3.**
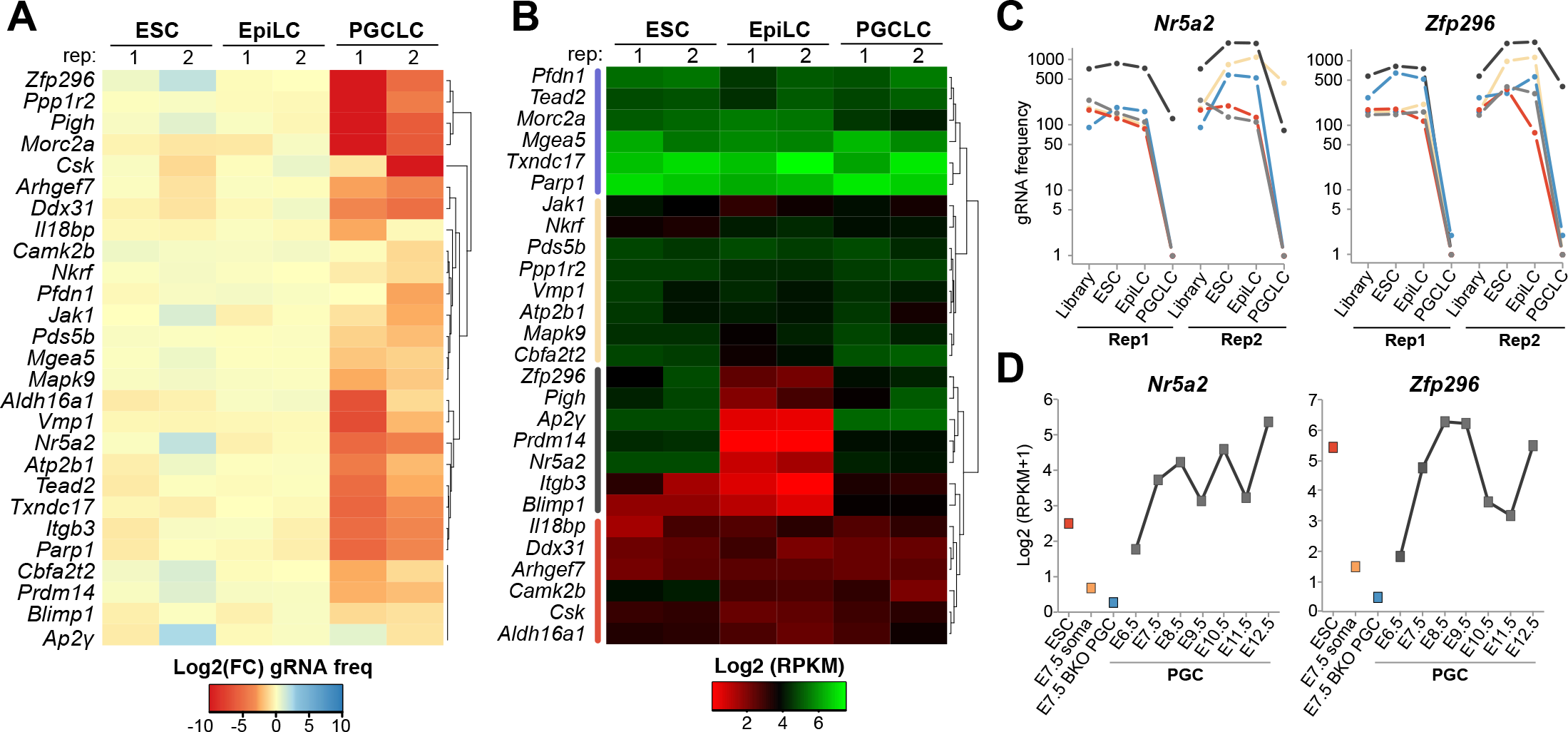
Candidate genes for primordial germ cell fate. **(A)** Heatmap showing mean fold-change (FC) in normalised frequency of gRNAs that target candidate PGC regulators, relative to their frequency in the preceding cell state. **(B)** Log_2_ RPKM gene expression dynamics of candidate PGC regulators during induction of SGET ESC into PGCLC in the screen. **(C)** Normalised frequency of individual gRNAs targeting *Nr5a2* and *Zfp296* during induction of PGCLC. **(D)** Expression dynamics of *Nr5a2* and *Zfp296* during *in vivo* formation of PGCs by single-cell RNA-seq. *Blimp1*^−/−^ PGC (BKO) and soma are shown for reference.

To refine our candidate list we examined their expression profile during induction of PGCLC from the transduced SGET ESC by RNA-seq. This confirmed that SG^+^ET^low^ PGCLC activate high levels of key PGC markers such as *Blimp1*, *Stella*, *Itgb3*, *Nanos3* and *Prdm14*, supporting their authentic germline identity (Fig S4D). The dynamics of candidate gene expression revealed four broad clusters, with cluster 1 and 4 corresponding to genes that are expressed relatively stably throughout ESC to PGCLC transition, with the latter nonetheless only just above the detection threshold implying it may filter out false-positives. In contrast, cluster 2 and 3 reflected dynamically regulated genes that are activated in PGCLC, with cluster 3 corresponding to genes fully silenced in EpiLC before strong reactivation, and including known PGC specifiers (Fig 3B). From these, we chose to focus on *Zfp296* and *Nr5a2*, since they rank 3 and 7 overall in the screen, and exhibit comparable PGCLC expression dynamics to *Prdm14* and *Blimp1* (Fig 3B & S4D). *Nr5a2* encodes an orphan nuclear receptor previously associated with reprogramming towards pluripotency, and also has a role in extraembryonic development, which leads to embryonic lethality when abrogated^25^. *Zfp296* encodes a zinc-finger protein that was amongst the original twenty Yamanaka factors, and has recently been shown to influence heterochromatin and iPS generation^26–28^.

Inspection of the individual gRNAs that target *Nr5a2* and *Zfp296* revealed a strong and reproducible reduction of their frequency specifically in PGCLC, but not in the preceding developmental transitions (Fig 3C). Moreover, single-cell RNA-seq indicates both genes are highly upregulated in nascent PGC *in vivo* at the very onset of specification, but not in adjacent somatic cells (Fig 3D). Both *Nr5a2* and *Zfp296* are also strongly upregulated in early human PGCs (Fig S4E), and fail to be activated in *Blimp1*^−/−^ PGC that are destined to diverge from the germline lineage (Fig 3D). The striking upregulation of these genes during incipient PGC specification, coupled with their high rank in the functional screen, prompted us to investigate their role in detail.

### Nr5a2 and Zfp296 regulate germline ontogeny

We used CRISPR editing to generate frameshifted null-alleles of *Nr5a2* or *Zfp296* in SGET ESC. *Nr5a2*^−/−^ ESC proliferated normally, expressed comparable levels of naïve pluripotency genes, and exhibited indistinguishable morphology from parental and matched WT ESC derived from the same targeting process. However upon induction of PGCLC, we observed a marked reduction in specification efficiency of *Nr5a2*^−/−^ cells at day 2 (d2) relative to WT controls (WT 19.8% ± 8.9; KO 6.4% ± 2.3) (Fig 4A). The impaired induction of Nr5a2-mutant cells was further reflected in day 6 (d6) PGCLC, and across multiple independent knockout lines (WT 44.7% ± 4.0; KO 17.4% ± 4.6), suggesting an important role for *Nr5a2* in PGC specification (Fig 4A & S5A). We next tested *Zfp296*^−/−^ ESC lines, which also exhibited normal morphology and pluripotent gene expression profiles, albeit with modestly reduced proliferation rate. Upon induction, PGCLCs were specified from *Zfp296*-mutant cells at apparently normal efficiency at d2 (WT 14.9% ± 3.2; KO 16.2% ± 2.2), as judged by the SGET reporter. However, by d6 we observed a highly significant depletion of PGCLCs in multiple mutant lines (WT 43.1% ± 3.5; KO 8.9% ± 2.8), suggesting the effects of *Zfp296* abrogation on PGCLC are progressive and manifest after specification (Fig 4B). More generally, these data indicate that loss-of-function of either *Nr5a2* or *Zfp296* leads to a significant impairment of germ cell development.

**Figure 4.**
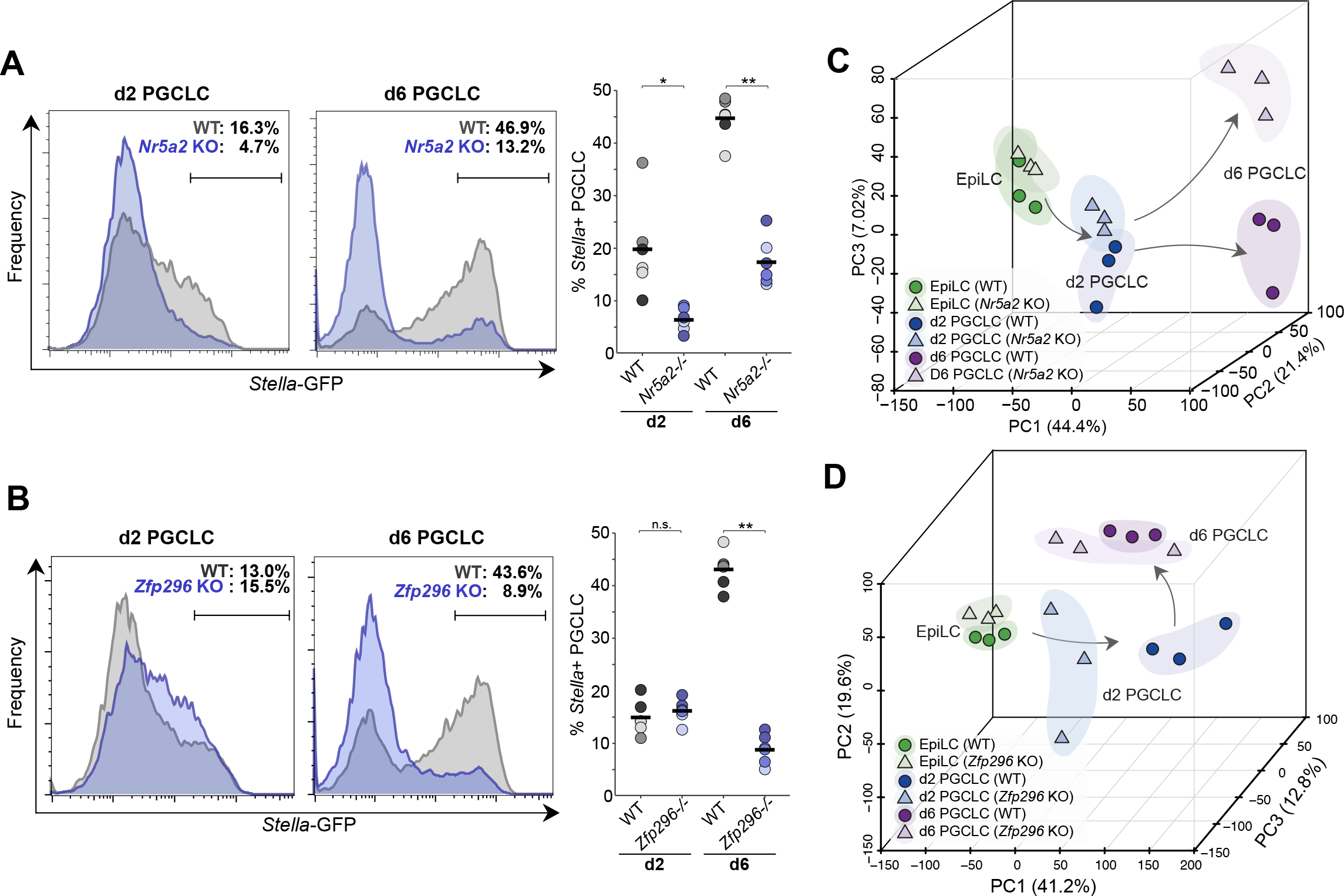
*Nr5a2* and *Zfp296* are key regulators of germ cell induction. **(A)** Representative FACS plot showing impaired induction of PGCLC in direct *Nr5a2*-knockout (KO) cells. Shown right is replicate quantifications of independent WT and KO lines. Independent lines are colour-coded. Bar indicates median value. **(B)** Representative FACS of PGCLC in *Zfp296*-knockout SGETs, with replicate quantification shown right. **(C)** Principal component analysis (PCA) showing the developmental trajectory of independent *Nr5a2*^−/−^ and matched-WT SGET lines during induction of PGCLC, based on global transcriptome **(D)** PCA of independent *Zfp296*^−/−^ and matched-WT SGET transcriptomes. * *p*<*0.05*; ** *p*<*0.01*.

Despite the dramatic reduction in numbers, both mutant-lines give rise to some putative PGCLC. To examine the identity of these cells, and their developmental trajectory, we performed RNA-seq. Unbiased hierarchical clustering revealed that *Nr5a2*^−/−^ EpiLC were interspersed with WT, but that PGCLC were clustered according to genotype, implying transcriptional differences in the absence of *Nr5a2* primarily arise upon induction of germline fate (Fig S5). To investigate this further we used principal component analysis (PCA), which confirmed that mutant EpiLC cluster closely with WT but that *Nr5a2*-null PGCLC progressively diverge from WT controls during their development (Fig 4C). By day 6 a strong distinction is apparent. This is consistent with *Nr5a2*^−/−^ EpiLC being competent for the germline lineage but undergoing an abnormal developmental trajectory upon PGCLC specification, which coincides with the timing of *Nr5a2* upregulation in WT cells.

Global transcriptome clustering of *Zfp296-null* cells indicated that precursor EpiLC were distinct yet highly comparable between WT and knockout lines (Fig S5). Indeed, PCA showed *Zfp296*^−/−^ EpiLC cluster closely with WT. In contrast, d2 mutant PGCLC exhibited a significantly different state from WT counterparts, that is closer to precursor EpiLC and thus apparently impaired in its developmental path towards germline fate, which is consistent with the subsequent loss of PGCLC numbers (Fig 4D). The remaining *Zfp296*-mutant PGCLC at day 6 appeared to partially re-establish the expected global profile, albeit with greater transcriptome variance between lines than WT. This implies that despite widespread transcriptional mis-regulation, a minor proportion of mutant PGCLC can overcome the effects of *Zfp296* abrogation, and potentially give rise to putative germ cells (Fig 4D). Overall, these data suggest that loss of *Nr5a2* or *Zfp296* has a dramatic influence on PGC development.

### Nr5a2 and Zfp296 converge on reciprocal regulation of WNT

To investigate the mechanisms that impairs development of mutant PGCLC and underpin their aberrant transcriptomes in more depth, we identified differentially expressed genes (adjusted *p*<*0.05*; >2 FC) in EpiLC and PGCLC. This revealed that downregulation of naïve pluripotency genes and upregulation of postimplantation markers proceeded similarly between *Nr5a2*^−/−^ and WT EpiLC counterparts (Fig 5A). Moreover, there was appropriate upregulation of early PGC markers such as *Nanos3, Stella, Blimp1* and *Prdm14* in d2 *Nr5a2*^−/−^ PGCLC, that was at least equivalent to WT, albeit with a modest failure to upregulate *Nanog*. However, we noted d2 *Nr5a2*-null PGCLC exhibited a highly significant over-expression of WNT pathway genes and targets, including *Cdx2*, *T* and *Mixl1* (Fig 5A). This was further reflected in gene ontology (GO) analysis of differentially expressed genes at d2, which highlighted *Canonical WNT signalling pathway* (adjusted *p*=*0.0061*) and *Pattern specification process* (adjusted *p*=*0.014*). We confirmed hyperactivation of key WNT pathway genes *T* and *Wnt3* in early *Nr5a2*−/− PGCLC using qRT-PCR on independent replicates (Fig 5B). Strong over-expression of the master mesoderm regulator *T* is predicted to override its role in germ cell induction and divert cells toward a somatic mesendoderm programme. Indeed, consistent with this, the surviving *Nr5a2*^−/−^ PGCLC at day 6 exhibit acute activation of mesendoderm regulators such as *Tbx4*, *Hey1* and *Pou2f3* (Fig 5A). This collectively suggests that the absence of *Nr5a2* leads to aberrant transduction of WNT signalling in nascent PGCLC, which in turn leads to upregulation of mesendoderm master-regulators and consequently de-repression of a somatic programme. It is possible that the aberrant activation of somatic genes in the absence of *Nr5a2* is a contributing factor that drives the majority of prospective PGCLC out of the germline programme, leading to reduced efficiency of specification.

**Figure 5.**
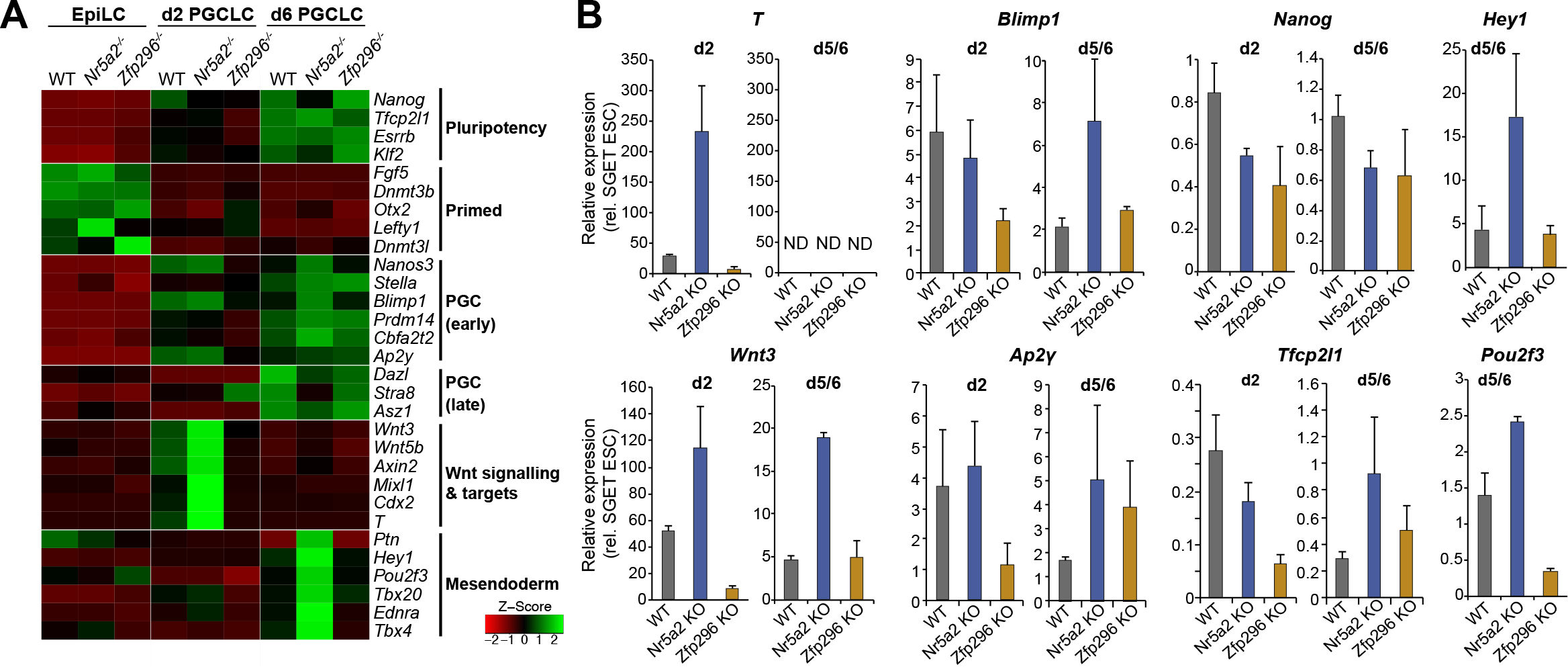
Compound mis-regulation of WNT and germ cell genes in the absence of *Nr5a2* and *Zfp296*. **(A)** Heatmap showing change in mean expression profile of selected genes from triplicate independent WT, *Nr5a2*^−/−^ and *Zfp296*^−/−^ lines during PGCLC induction by RNA-seq **(B)** Quantitative validation of gene expression changes in PGCLC from WT, *Nr5a2*^−/−^ and *Zfp296*^−/−^ lines by independent qRT-PCR experiments. Error bars show s.e.m of duplicate biological experiments, each comprising at least twelve independent replicate inductions. ND, not detected.

We next analysed gene expression in *Zfp296*-null EpiLC, which revealed that pluripotency genes including *Nanog, Esrrb* and *Tfcp2l1* were repressed as expected, whilst EpiLC markers such as *Fgf5* and *Dnmt3b* were appropriately upregulated. In contrast mutant PGCLC at day 2 exhibited a clear signature of mis-expressed genes (Fig 5A). Remarkably, GO analysis again suggested this primarily corresponded to WNT targets, but in in contrast to hyperactivation of WNT in *Nr5a2*^−/−^ PGCLC, *Zfp296-null* PGCLC exhibited a severe block of WNT-associated gene activation; the top enrichment is *Negative regulation of canonical WNT signalling* (adjusted *p*=*0.0064*). Amongst all genes *T* is the most significantly downregulated in early *Zfp296*-mutant PGCLC, with *Cdx2, Notum* and *Dkk1* also being amongst the top 20. The absence of appropriate level WNT signalling could manifest as impaired activation of critical germline genes that are WNT and/or *T* targets^5^. Indeed, *Zfp296*-null PGCLC at d2 exhibited a block in *Blimp1, Prdm14 Cbfa2t2a* and *Ap2γ* activation (Fig 5A). qRT-PCR analysis independently confirmed highly impaired *T* and WNT genes in *Zfp296*-mutant PGCLC and concomitant significant reduced activation of germline regulators *Blimp1* and *Ap2γ* (Fig 5B). This suggests that absence of *Zfp296* establishes a reciprocal developmental failure relative to deletion of *Nr5a2*, with both genes converging on regulation of appropriate WNT signalling in nascent PGCLC. In the case of *Zfp296*, this appears to manifest in a delayed but dramatic loss of PGCLC by d6. Notably, the SG'ET” cells from both *Nr5a2* and *Zfp296* embryoids, which have putatively acquired somatic fate, exhibited comparable expression of lineage-specific markers *Sox17, Hoxa1* and *Sox7* relative to wild-type (Fig S6A). This implies that the loss of function for both genes does not impede the exit from the pluripotent state towards somatic fates, but these cells are impaired specifically in their capacity to form PGCs.

### Rescue of Nr5a2 and Zfp296 deficiency

We considered that the impaired specification and aberrant transcriptome of mutant PGCs could be due to impaired epigenetic reprogramming, as reported in *Prdm14*^−/−^ PGC^10^. To investigate this, we quantified global DNA methylation erasure and observed that WT, *Nr5a2*^−/−^ and *Zfp296*^−/−^ PGCLC all underwent extensive and comparable DNA demethylation (Fig 6A). This was confirmed by bisulfite pyrosequencing analysis of LINE-1 elements (Fig S6B). We therefore turned our attention back to understanding whether the observed mis-regulation of WNT could be driving impaired germline fate, by exposing incipient PGCLC to WNT inhibitor (WNTin: XAV939) or WNT agonist (WNTag: Chiron). Addition of WNTin led to strong downregulation of key PGC genes including *T* and *Wnt3*, as well as *Nanog*, consistent with the effects of impaired WNT transduction in *Zfp296-null* PGCLC (Fig 6B). Reciprocally, WNTag affected an increase in expression of *T* and *Wnt3* at d2, whilst also downregulating *Nanog*, which is in line with the effects of WNT over-activity in *Nr5a2*-null PGCLC. Indeed, WNTag also elicited overexpression of mesendoderm gene *Hey1* by d6 PGCLC (Fig 6C). Notably WNTinh led to a significant depletion of specified PGCLC (*p*<*0.05*), whilst partial WNT activation by WNTag also trended towards impaired PGCLC specification (Fig 6D). These data suggest that precise levels of WNT activity are necessary for optimal PGC specification.

**Figure 6.**
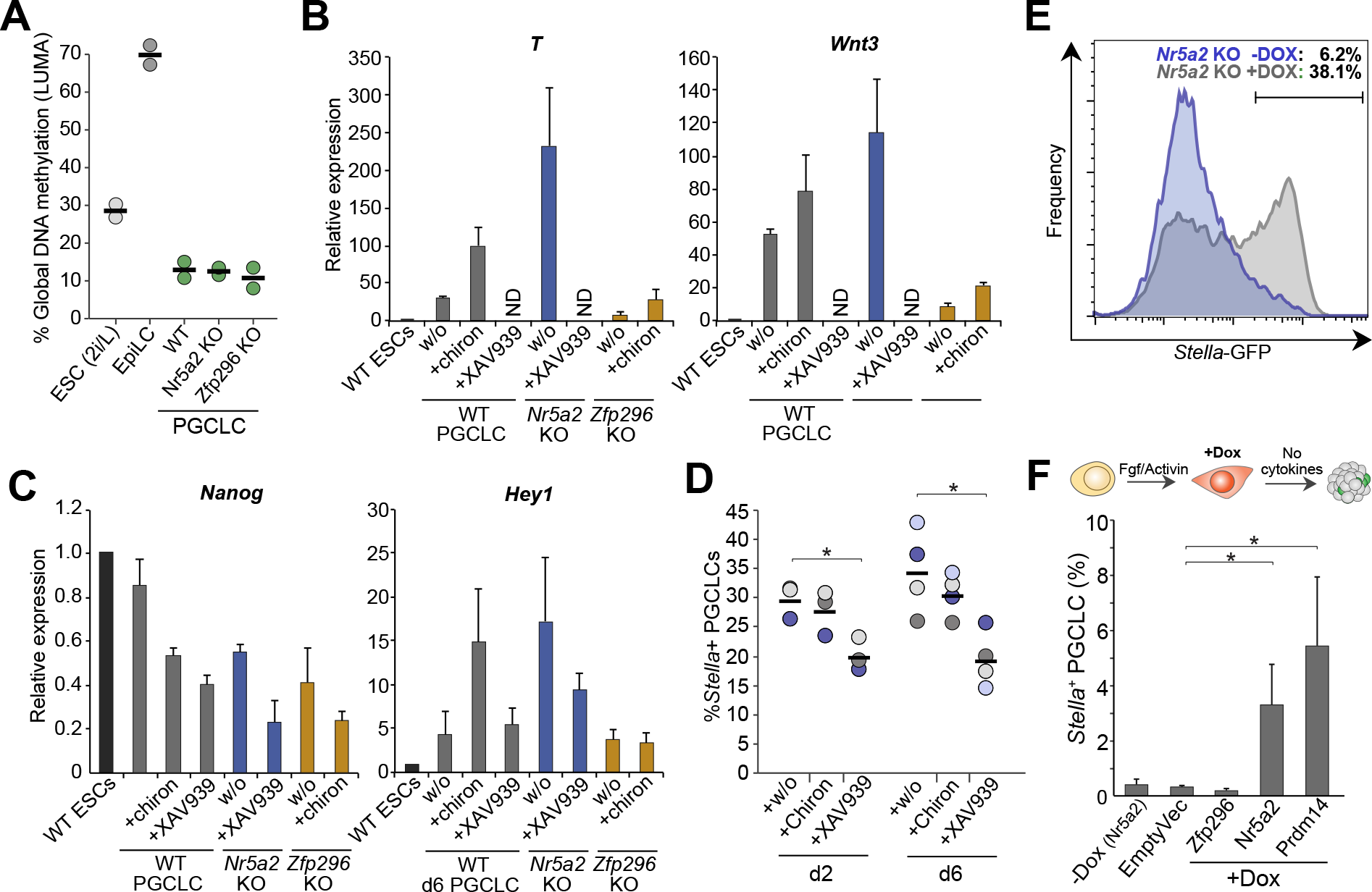
Role of WNT and rescue of *Nr5a2* defects. **(A)** Global levels of DNA methylation reprogramming in WT and mutant PGCLC assayed by LUMA. **(B)** qRT-PCR analysis showing expression of WNT genes and targets are affected similarly to knockouts by WNT inhibitor (XAV939) or agonist (Chiron). **(C)** Expression of *Nanog* and mesendoderm gene *Hey1* by qRT-PCR in WT and mutant PGCLC with WNT modulation. **(D)** Effects of WNT manipulation on efficiency of PGCLC ontogeny in quadruplicate independent experiments. Bar indicates median value and *=*p*<*0.05*. **(E)** FACS plot showing the *Nr5a2*-mutant PGCLC specification defect at d2 is rescued by Dox-inducible activation of exogenous *Nr5a2* cDNA. Dox is added after EpiLC induction. **(F)** Percentage *Stella*^+^ PGCLC induced without cytokines from WT cells upon DOX-inducible expression of the indicated cDNA.

Addition of WNTin to *Nr5a2*^−/−^ PGCLC rescued hyperactivated *T* and *Wnt3*, demonstrating that their upregulation in *Nr5a2-*-mutant PGCLC is likely a direct consequence of de-regulated WNT activity (Fig 6B). In contrast addition of excess WNTag to *Zfp296*-mutant PGCLC only resulted in modest rescue of aberrantly repressed WNT targets, suggesting that in the absence of *Zfp296*, germ cells cannot respond to WNT activation, even when forced. We therefore attempted to rescue the specification defects by generating mutant PGCLC that carry Doxycycline(DOX)-inducible expression of wild-type *Nr5a2* and *Zfp296*. We induced *Nr5a2* expression specifically during PGCLC induction in a *Nr5a2*^−/−^ background, which led to a highly significant rescue of PGCLC specification. Moreover, this reimposed supression of somatic and WNT target genes, implying a direct effect of *Nr5a2* on these pathways (Fig 6E & S6C-D). In contrast, we found expression of *Zfp296* to be highly toxic to both ESC and PGCLC, even at very low background (leaky) levels, suggesting precise control over its expression level is important.

Remarkably DOX-induced *Nr5a2* expression in mutant PGCLC resulted in increased PGCLC induction as compared to wild-type counterparts. This encouraged us to test whether *Nr5a2* could drive germ line fate independently of BMP cytokines that are normally requisite. Here, forced expression of *Nr5a2* in wild-type EpiLC led to emergence of SG^+^ET^low^ PGCLC at >10-fold higher efficiency than background; at levels comparable with forced expression of canonical PGC-specifier *Prdm14*, albeit at low absolute frequency (3.3%) (Fig 6F). Induction of *Nr5a2* resulted in PGCLC with apparently similar expression profiles as authentic PGCs (Fig S6E). This indicates that activation of *Nr5a2* can, at low efficiency, directly induce PGC-like fate from competent EpiLC. More generally, our data places *Nr5a2* and *Zfp296* as critical regulators that provide robustness to germline specification by modulating appropriate WNT activity.

## DISCUSSION

Our study outlines a principle for using CRISPR screens to deconvolve the genetic basis of complex cell fate decisions. By establishing a ‘traffic-light’ SGET reporter system we traced the pathway from pluripotency to germ cell fate, and identified important genes for each transition. This strategy can be transposed to interrogate multi-step transitions towards any lineage for which a faithful compound-reporter or isolation method can be developed.

Utilising SGET we identified genes involved in exiting naïve pluripotency, including the previously described key regulators *Zfp281* and *Tcf3*^29,30^, and further validated roles for novel candidates *Zmym2*, *FoxP1* and *Uchl5*. Amongst these *Zmym2* expression is anti-correlated with naïve pluripotency genes in heterogeneous single-ESC^31^, and is upregulated when cells are induced to egress from naïve pluripotency both *in vitro* and *in vivo* (Fig S3). Indeed, ZMYM2 is known to directly interact with NANOG^32^, and its negative regulatory activity could potentially blunt NANOG function thereby promoting exit from pluripotency, or it could alternatively function in this role as part of the LSD1 complex^33^. Interestingly the transcription regulator *FoxP1* is also strongly activated during EpiLC induction and, whilst it can be alternatively spliced to generate a pro-pluripotency isoform^34^, our data implies canonical *Foxp1* isoforms play a role in antagonising naïve pluripotency. The potential mechanism of function for the deubiquitinase *Uchl5* is less clear, but it has previously been associated with regulation of the TGF-β/Smad pathway^35^. Future work will be important to characterise the regulatory role of the full repertoire of candidates in control of pluripotent status.

We further exploited the SGET strategy to identify *Zfp296* and *Nr5a2* as key functional genes for the germline lineage and as important parts of an expanded PGC gene regulatory network. Deletion of either *Nr5a2* or *Zfp296* converges on mis-regulation of the WNT pathway, resulting in its aberrant activation or inhibition respectively, implying that precisely regulated levels of WNT is fundamental for germ cell ontogeny. Indeed, loss of either gene leads to severely disrupted PGCLC development. Of note *Nr5a2* is a reported WNT target^36^ and directly interacts with β-catenin^37^, raising the possibility it forms a negative-feedback loop with WNT to modulate appropriate signalling levels in PGCs. This must strike a balance between WNT-mediated activation of germ cell regulators such *Blimp1* and *Prdm14 ^5^*, and the capacity to subsequently repress the WNT co-activated mesodermal programme. By blunting WNT activity, NR5A2 may ensure the threshold is tipped towards reversibility, since in its absence PGCLC development is impaired and linked with aberrant activation of mesendodermal genes. Functionally, *Nr5a2* was previously shown to be important for reprogramming somatic cells to iPS cells but not for pluripotency *per se* ^38–40^. Similarly, PGC undergo reprogramming but do not acquire pluripotency^4^, supporting a role for *Nr5a2* in resetting PGCs away from somatic fates. Notably, NR5A2 binding motifs are among the most significantly enriched within germline-associated promoters, suggesting *Nr5a2* could also have a direct function on PGC genes^41^.

In contrast to *Nr5a2*, loss of *Zfp296* results in failure to appropriately upregulate WNT in the germline, as well as affecting some TGF-β targets such as *Lefty1*. Underactive WNT in *Zfp296*-mutant PGCLC manifests as impaired activation of early germline genes, including *Blimp1* and *Prdm14*. This in turn leads to dramatic loss of PGCLC. Indeed, a recent report has shown the absence of ZFP296 leads to sterility, confirming the long-term consequences of its loss-of-function on the germline^27^. The molecular mechanism through which *Zfp296* impacts WNT to regulate PGC ontogeny is an important topic for future research. ZFP296 can however influence global H3K9 methylation and has been reported to negatively regulate KLF4 activity^27,42^, which counteracts murine *Blimp1* expression^43^, implying *Zfp296* could contribute to the PGC programme through multiple regulatory routes. Of note, both *Zfp296* and *Nr5a2* are strongly activated in nascent human PGC (Fig S4), comparably to their upregulation in mouse PGC, indicating they may have a broad conserved role in mammalian germline development.

Our approach and experimental design is both objective and based on functional outcomes, thereby delineating an alternative strategy to identify critical components of cell-type specific gene regulatory networks, which in the case of germ cells, has previously been challenging. The approach preferentially identifies intrinsic cell fate regulators however; not weak-acting or extrinsic regulators, for at least two reasons. Firstly, the addition of extrinsic signalling at high levels (FGF and ACTIVIN for EpiLC; BMP among others for PGCLC) overcomes weak-or signalling receptor-specific effects by acting on multiple redundant pathways to drive fate transitions. Second, the pooled screen strategy means that knockout cells for a given extrinsically-acting gene (e.g. *Wnt3*) can be rescued by paracrine effects from adjacent cells with wildtype alleles. For these reasons, FGF signalling and WNT components were not identified in the exit from pluripotency and PGCLC screens, respectively. Nevertheless, we identified multiple cell-intrinsic candidates. Indeed, recent optimisations to loss-of-function CRISPR libraries^44^ will minimise false-negatives in future iterations, which can arise due to ineffective gRNAs confounding significance scores, as we observe for *Prdm14* for example. In summary, we have used CRISPR screening to identify key regulators for successive cell fate transitions out of naïve pluripotency towards the germ cell lineage. In doing so we characterised *Zfp296* and *Nr5a2* as crucial for driving germ cell development by acting to regulate appropriate WNT levels, thereby controlling the balance between activation of PGC genes and silencing of the somatic programme.

## AUTHOUR CONTRIBUTIONS

J.A.H designed the study, performed experiments and bioinformatics, and wrote the manuscript. Y.H performed experiments and bioinformatics. U.G performed experiments and wrote the manuscript. K.H.G performed support experiments. T.K performed experiments. M.A.S supervised the study and wrote the manuscript.

## ACKNOWLEDGEMENTS

Funding for this study came from a Wellcome Trust programme grant and Cancer Research UK (C6946/A14492) / Wellcome Trust (092096) core grants to M.A.S. Further funding came from a core European Molecular Biology Laboratory (EMBL) grant to J.A.H.

## COMPETING FINANACIAL STATEMENT

We declare no competing financial interest.

## MATERIALS & METHODS

### ESC culture and PGCLC induction

ESCs were maintained in N2B27 medium supplemented with 2i (PD0325901 (1 μM) and CHIR99021 (3 μM); both Stemgent), 1000 U/ml LIF (Cambridge Centre for Stem Cell Research) (2i/L) and 1% Knockout Serum Replacement (KSR) on fibronectin at 37°C in a 5?% CO_2_ humidified chamber. For serum/LIF (S/L) culture, cells were maintained on gelatin in GMEM supplemented with 10% FCS, L-Glutamine, non-essential amino acids (NEAA), 0.1 mM β-mercaptoethanol and 1000 U/ml LIF. Induction of EpiLC and PGCLC was carried out as previously described (Hayashi et al., 2011; Hayashi and Saitou, 2013). Briefly, ESCs cultured in 2i/L were washed thrice with PBS, and seeded onto fibronectin (16.7 mg/ml) coated plates with EpiLC medium (N2B27, 1% KSR, bFGF (12 ng/ml), Activin A (20 ng/ml)) for 42hrs. Media was changed every day. EpiLCs were then gently dissociated and seeded at 3000 cells per embryoid in ultra low-cell binding U-bottom 96-well plates (NUNC) with PGCLC induction medium (GK15: GMEM, 15% KSR, NEAA, 1 mM sodium pyruvate, 0.1 mM β-mercaptoethanol, 100 U/ml penicillin, 0.1 mg/ml streptomycin, and 2 mM L-glutamine; supplemented with cytokines: BMP4 (500 ng/ml), LIF (1000 U/ml), SCF (100 ng/ml), BMP8a (500 ng/ml), and EGF (50 ng/ml)). For large scale (CRISPR screen) PGCLC inductions (>4,000 embryoids per replicate), 30 μl of PGCLC medium was used for each embryoid, with 20μl additional PGCLC medium added on day 2. For smaller-scale PGCLC inductions, 150 μl of PGCLC medium was used per embryoid and another 50μl of PGCLC medium was added on day 2. To minimise evaporation, only the inner 60 wells were used during the screen and 200μl of PBS was added to the outer 36 wells. After 2-6 days, EBs were collected for fluorescence-activated cell sorting (FACS) or gene expression analyses. SGET EpiLCs were also seeded in GK15 medium without cytokines as a negative control to normalise for background levels of SGET fluorescence in embryoids. Where indicated, WNT inhibitor XAV939 (10 μM) or agonists CHIR99021 (3μM) were added upon induction of PGCLC in embryoids. Where indicated doxycycline was added at 200 ng/ul upon induction of PGCLC in embryoids +/− above mentioned cytokines.

### Generation and validation of the SGET reporter

Embryonic stem cell lines carrying *Stella*-GFP BAC (SG)^45^ were derived as previously described, and routinely checked for mycoplasma-negative status using the mycoalert detection kit (Lonza). These lines were used as the substrate to target an in-frame tdTomato cassette into exon 1 of *Esg1* (ET) by homologous recombination, thereby generating SGET ESC. Targeting vector(s) for *Esg1* were constructed by amplifying homology arms from genomic DNA of mouse ESCs by PCR using PrimeSTAR MAX (Takara Bio, Otsu, Japan), tdTomato from ptdTomato-N1 (Takara Bio, Otsu, Japan) anda destabilized domain (d2) with T2A BSD from pL1L2_IRESdCherryT2Abla-pAneotk vector (a gift from core facility in Cambridge Stem Cell Institute), and assembled using InFusion HD cloning kit (Takara Bio, Otsu, Japan). For gene targeting, 1 × 10^7^ low-passage (<10) *Stella*-GFP ESC suspended in PBS were electroplated with 40 μg linearized targeting vector using conditions: 800V, 10F 49 in Gene Pulser equipment (Bio-Rad). ESCs were selected with 10 mg/ml blasticidin, and colonies were picked and screened for correct targeting by PCR. We tested for faithful reporter activity using tetraploid embryo complementation (~E8.5) and chimera formation assays (E12.5). For tetraploid complementation assay, 2-cell stage diploid embryos were collected in M2 medium from the oviduct of mice at E1.5, and washed 3 times with medium containing 0.01% polyvinyl alcohol (Sigma), 280 mM Mannitol (Sigma), 0.5 mM Hepes (Sigma), and 0.15 mM MgSO4 (Sigma). Electrofusion of blastomeres to produce tetraploid embryos was subsequently carried out using a DC pulse (100 V/mm, 30 msec, 1 time) followed by application of AC pulses (5 V/mm, 10 sec) using ECM 200 (BTX, Holliston, MA). Tetraploid embryos were transferred into KSOM medium (Merck Millipore) and cultured for 24 hours for 4-cell / morula injection. For chimera formation assay, morula stage diploid embryos were collected in M2 medium from the oviduct of mice at E2.5. For micro-manipulation, a piezo-driven micro-manipulator (Prime Tech, Tokyo, Japan) was used to drill through the zona pellucida and 5-10 SGET ESCs were introduced into the perivitelline space of morula stage tetraploid or diploid embryos. They were cultured to the blastocyst stage and then transferred into the uteri of pseudopregnant recipient MF1 female mice (E2.5). Post- implantation embryos were dissected at ~E4.5, E6.0, E7.0, E8.5 and E12.5 to analyse eGFP and tdTomato expression.

### Lentiviral CRISPR screen

A genome-wide lentiviral CRISPR gRNA library was utilised that contains 87,897 gRNAs targeting 19,150 mouse protein-coding genes (Yusa et al), with up to 5 gRNAs per gene (Addgene: #50947). The gRNA library was amplified with NEB 10-beta electrocompetent *E.coli* (NEB) as per the recommended protocol. Briefly, *E.coli* were transformed via high efficiency electroporation, to ensure faithful library replication, and incubated at 37°C for 1 hour with SOC recovery medium (ThermoFisher) before growing in 500ml 2xTY (16 g/L Tryptone, 10 g/L Yeast Extract, 5.0 g/L NaCl) + ampicillin (50 μg/ml) mediums at 37°C overnight with 230 rpm shaking. Plasmid was purified from 500 ml bacterial cultures using an endotoxin-free plasmid maxi kit (Qiagen) as per manufacturer’s instructions. Faithful replication/amplification of the library was confirmed by Illumina sequencing. For lentiviral production, 4×10^6^ HEK 293T cells cultured in DMEM (Invitrogen), 10% FCS, L-glutamine and Penicillin-Streptomycin were seeded onto 10 cm poly-lysine coated plates. The following reagents were prepared in OptiMEM per plate: 10 μg CRISPR plasmid library, 10 μg VSVG-pcDNA3 (Addgene:#8454), 10 μg pCMV-dR8.2 (Addgene:#8455) and 30 μl of Lipofectamine 2000 (ThermoFisher Scientific), and incubated with cells for 5-6 hours at 37°C in a 5% CO_2_ incubator. Medium was changed to DMEM with Forskolin (5 μM/ml) and viral supernatant was collected after 48 hours and 64 hours. Cell debris was removed by centrifugation at 1200 rpm for 15 minutes at 4°C, and the supernatants were filtered with 0.45 μM filter. To concentrate the virus supernatant, it was combined with 4xPEG solution (320ml of 50% PEG 6000, 40 ml of 5 M NaCl, 20 ml of 1 M HEPES, adjust pH to 7.4 and add ddH20 until 500 ml). The virus/PEG mixture was incubated for 4 hours at 4°C, centrifuged at 2600g for 30 minutes at 4°C, and the supernatant aspirated; the process was repeated twice. The viral titre (multiplicity of infection) was assayed by determining the percentage of cells with viral integration after transduction by flow analysing BFP-positive cells and confirmed by qPCR (for BFP).

For the CRISPR screen we first generated SGET ESC that carried constitutive expression of *Cas9* by introducing a CAG-*Cas9*-pA cassette into SGET cells via piggybac-mediated transposition. Clones were subsequently picked and screened to identify a single-copy integration of *Cas9*, and to confirm functional capacity to direct high-efficiency gRNA-directed DNA indels using a GFP reporter. To introduce the CRISPR library, a total of 5×10^7^ SGET-*Cas9* ESC were transduced with lentiviral library using a multiplicity of infection (MOI) of 0.3, with biological replicates performed on independent occasions. Transduced SGET ESC were maintained on fibronectin-coated T175 flasks in N2B27, 2i/LIF and 1% KSR, with 8ng/μl of polybrene added during lentiviral infection. Medium was changed 24 hours later and cells were selected with puromycin (1.2μg/ml) for 5 days. High numbers of transduced ESC were passaged every 2-3 days at low ratio to ensure the complexity of the gRNA library was maintained. ESC pellets were taken at day 12 for analysis and EpiLC were induced on day 12 post-lentiviral transduction, from which PGCLC were subsequently induced.

To quantify the relative frequencies of integrated gRNAs in each population during cell fate transitions, genomic DNA was isolated from day 12 ESC post-lentiviral transduction, from SG^−^ET^+^ and SG^+^ET^+^ EpiLC populations at 42hr, and from SG^+^ET^low^ PGCLCs from day 6 embryoids, in biological duplicate using the DNeasy Blood & Tissue kit (Qiagen). Purified genomic DNA was used as template for custom primers that specifically amplify the gRNA region, and include overhanging Illumina adaptors and indexes to allow deep sequencing and multiplexing, respectively (synthesised as Ultramers from IDT) (see oligo tables). Prior to PCR, genomic DNA was sonicated with a Biorupter (Diagenode) for 15 seconds on ‘LOW’ power to improve efficiency of amplification. gRNA sequences were amplified in multiple PCR reactions to enable all isolated DNA to be utilised, each with NEBNext^®^ Q5 HotStart HiFi PCR Master Mix (NEB), 0.2 μM of universal forward primer (mix of staggers), 0.2 μM of indexed reverse primer and genomic DNA (~650 ng / 50 *μ*l rxn). For amplification from the plasmid vector, 15 ng was used as template for replicate amplifications. The following cycling conditions were used: 95°C for 2 minutes, then between 21-26 cycles of (98°C for 20 seconds, 62°C for 20 seconds and 72°C for 20 seconds) followed by 72°C for 1 minute and hold at 4°C. The number of cycles was optimised for the point at which PCR products can be first visualised on an agarose gel. The PCR reaction was subsequently purified with AMPure^®^ XP beads (Beckman Coulter) using double size selection to remove primer dimers and genomic DNA. Briefly, 0. 55x of AMPure beads were added to PCR reaction (1x) and incubated for 5 minutes at room temperature. The supernatant was collected; the beads contained the unwanted larger fragments and were discarded. An additional 0.3x of AMPure beads was subsequently added to the supernatant (1x) and incubated for 5 minutes at room temperature. The supernatant was removed and beads washed twice with 200*μ*l of 80% EtOH, air dried for 5 minutes and DNA was eluted from AMPure beads with EDTA-low TE buffer. The concentration of adaptor ligated gRNA amplicons was measured with the Qubit DNA Assay kit (ThermoFisher Scientific) and the fragment distribution determined with an Agilent D1000 ScreenTape System (Agilent Technologies). Libraries were subsequently multiplexed and sequenced with an Illumina HiSeq 1500 using single-end 50bp reads. As the sequences for all samples were identical up to the gRNA region, these types of low complexity libraries can produce low quality data. To counteract this, forward staggered primers were introduced at equal ratios during PCR amplification, generating offset reads. In addition, libraries were sequenced with 4 dark cycles and low density (70%) clustering.

### Gene editing in ESC cell lines

Targeted knockout ESC lines were generated using CRISPR genome editing by either deleting a critical exon to generate a frame-shifted null-allele or via inducing frame-shifting null indels into early coding exonic sequences. In brief, 20nt gRNAs encoding complementary sequences to the region/gene to be targeted were cloned into px459 (v2.0) (Addgene:#62988) using the *Bbs1* sites and transfected into SGET ESC with lipofectamine 2000, according to the manufacturers guidelines. A table of the gene targeting gRNAs and the editing strategy is shown below. Transfection of ESC was selected for with 1.2 ug/ml puromycin for 48 hours, and ESC were subsequently passaged into 6-well plates at low density (1,000-5,000 ESC per well). One week later colonies were picked and expanded before undergoing genotyping by amplicon sizing and sanger sequencing to determine the precise mutant sequence at each allele using the tracking of indels by decomposition tool (https://tide.nki.nl). Multiple mutant lines were catalogued and validated before use in functional assays, where WT and/or heterozygous control clones derived from the same targeting process were utilised. For doxycycline inducible expression, *Zfp296, Nr5a2* or *Prdm14* cDNA was cloned into a custom PiggyBac vector downstream of a doxycycline-responsive promoter, and transfected into SGET ESC in conjunction with the reverse transactivator (TET-3G). Genomic integration was selected for with 250ug/ml neomycin.

### Flow cytometry

For fluorescence-activated cell sorting (FACS), cultured cells or embryoids were dissociated into single cells with TryLE, and suspended in phosphate buffered saline (PBS) with 0.5% BSA or 3% FBS. Filtered cell suspensions were subsequently sorted using a Sony SH800Z or a MoFlow high-speed cell sorter (Beckman Coulter) or analysed with BD LSRFortessa X-20 (BD Biosciences). For exit from pluripotency analyses, cells were sorted/analysed based on absence or presence of *Stella*-eGFP expression, whilst maintaining *Esg1*-tdTomato activitaion (SG+/−ET+). For PGCLC, cells were sorted/analysed based on *Stella*-eGFP re-activation coupled with low *Esg1*-tdTomato. The threshold(s) levels for sorting was set and normalised based on expression levels in pre-optimised SGET ESC. A negative population of ESCs without any fluorescence was used to set absolute thresholds. Forward and side scatters were used to gate for the cell population and doublets. For each sort the maximum number of cells was collected, whilst for analysis at least 10,000 cell datapoints was captured to achieve a representative sample.

### RNA-seq and Gene Expression

For RNA-seq, EpiLC, day 2 or day 6 PGCLC were dissociated into single cell solutions with TrypLE and sorted with a Sony SH800Z or a MoFlow high-speed cell sorter (Beckman Coulter) based on appropriate *Stella*-eGFP and *Esg1*-tdTomato expression. Samples were sorted into 150μl of extraction buffer from the PicoPure RNA isolation kit (Life Technologies) and rapidly frozen on dry ice or liquid nitrogen. Total RNA was subsequently extracted using the PicoPure RNA isolation Kit, including a 15 minute on-column DNaseI digestion. RNA integrity number was assessed with RNA HS? ScreenTape (Agilent), and all samples confirmed to have a RIN >8.5. For RNA-seq from CRSIPR screen samples, 50 ng of total RNA was used as input for the Ovation RNA-seq System v2 (Nugen) as per manufacturer’s instructions. Amplified double-stranded (ds) cDNA was diluted into EDTA-low TE (Agilent) and sheared into ~230bp in length using S220 Focused-Ultrasonicator (Covaris) using settings: duty factor 10%, cycle burst 200, intensity 5, temp at 4°C and treatment time of 5 minutes per sample. A high Sensitivity D1000 ScreenTape assay (Agilent) was used to assess the efficiency of library fragmentation. Fragmented ds-cDNA was concentrated with Qiagen Reaction Clean Up kit (MiniElute) and 1 μg of the fragmented ds-cDNA used as input for the library preparation using Encore Rapid DR Multiplex Library System (Nugen). This kit ligated the adaptors to repaired-end ds-cDNA without amplification, which eliminated biases introduced during PCR amplification. The KAPA Library Quantification Kit (Kapa bioscience) was used to quantify the concentration of each adaptor-ligated libraries prior to multiplexing. The ESC, EpiLC, PGCLC and soma cell RNA sequencing libraries were generated in parallel for each replicate, and the biological replicates were generated on independent occasions. Samples were multiplexed and sequenced with Illumina HiSeq1500, single-end 50bp read length with a minimum depth of approximately 15 million reads per sample.

For RNA-seq of *Nr5a2*^−/−^, *Zfp296*^−/−^ and matched WT control cell lines, 100 ng of total RNA from triplicate biological independent experiments was used as inputs for the NEBNext Ultra RNA library Prep Kit for Illumina^®^ (NEB), with the library generated as per manufacturer’s instructions. During the final PCR amplification stage, 15 cycles of amplifications were performed to generate the adaptor ligated, fragmented cDNA for sequencing. Samples were assessed using the High Sensitivity D1000 ScreenTape assay (Agilent) to ensure the library did not contain primer-dimer contamination after last round of AMPure beads cleaning. The Qubit dsDNA HS assay kit (ThermoFisher Scientific) and the NEBNext Library Quant Kit for Illumina^®^ was used to accurately quantify the concentration of each library preparation. Samples were multiplexed with 12 indexes per lane and a total of 2 lanes of sequencing on Illumina HiSeq4000, single-end 50bp with an average depth of approximately 20 million reads per sample.

### DNA methylation

Global DNA methylation levels were determined using the LUminometric Methylation Analysis (LUMA) method^46^ and bisulfite pyrosequencing. Briefly, genomic DNA was isolated from purified PGCLC using the DNeasy Blood & Tissue kit (Qiagen) and treated with RNase. 50-100ng of DNA was digested with MspI/EcoRI and HpaII/EcoRI (NEB) and the subsequent methylation-sensitive overhangs were quantified by Pyrosequencing (PyroMark Q24 Advance) with the dispensation order: GTGTGTCACACAGTGTGT. Global CpG methylation levels were determined from relative peak heights at position(s) 7, 8, 13 and 14 using the formula: [(2*(p7*p13))/(p8+p14)]^HpaII^/[(2*(p7*p13))/(p8+p14)]^MspI^. CpG methylation at LINE1 loci was determined by bisulfite pyrosequening using the EpiTect bisulfite kit (Qiagen). PCR amplification and assay design were performed as previously described.

### Bioinformatics

For RNA-seq, expression reads were quality-trimmed using *TrimGalore* to remove adapters, and aligned to the mouse reference genome (GRCm38/mm10) using *TopHat2* guided by ENSEMBL gene models. Mapped reads were imported into *SeqMonk* analysis software and quantitated using the RNA-seq quantitation pipeline. Differentially expressed genes (DEG) were identified using the DESeq2 statistical package with significance thresholds set as *p*<*0.05*, filtering for a fold-change (FC) > *2* and mean RPKM expression >1 in at least one normalised sample. Analysis of gene ontology enrichment was performed using DEG datasets in the DAVID bioinformatics resource v6.7 (https://david.ncifcrf.gov) and/or the REVIGO visualisation tool (http://revigo.irb.hr). Heatmaps and plots were generated using custom scripts in the R statistical package. For analysis of the CRISPR screen, demultiplexed gRNA sequences were extracted from Illumina reads using custom scripts to account for staggered start positions. These were quantitated and statistically analysed using Model-based Analysis of Genome-wide CRISPR-Cas9 Knockout (MAGeCK) v.0.5.2 software in python^47^. Library sequences were trimmed to exclude the bottom 1% of reads from the initial library, which primarily corresponded to zero-count gRNAs. MAGeCK analysis was performed using a ‘total’ normalisation method to compare gRNA counts with the preceding cell population during ESC-EpiLC-PGCLC transition (or with the vector library in the case of ESC), taking genes with *p*<*0.01* for negative- or positive- enrichment in independent replicate screens as candidates.

### Oligos

#### qRT-PCR

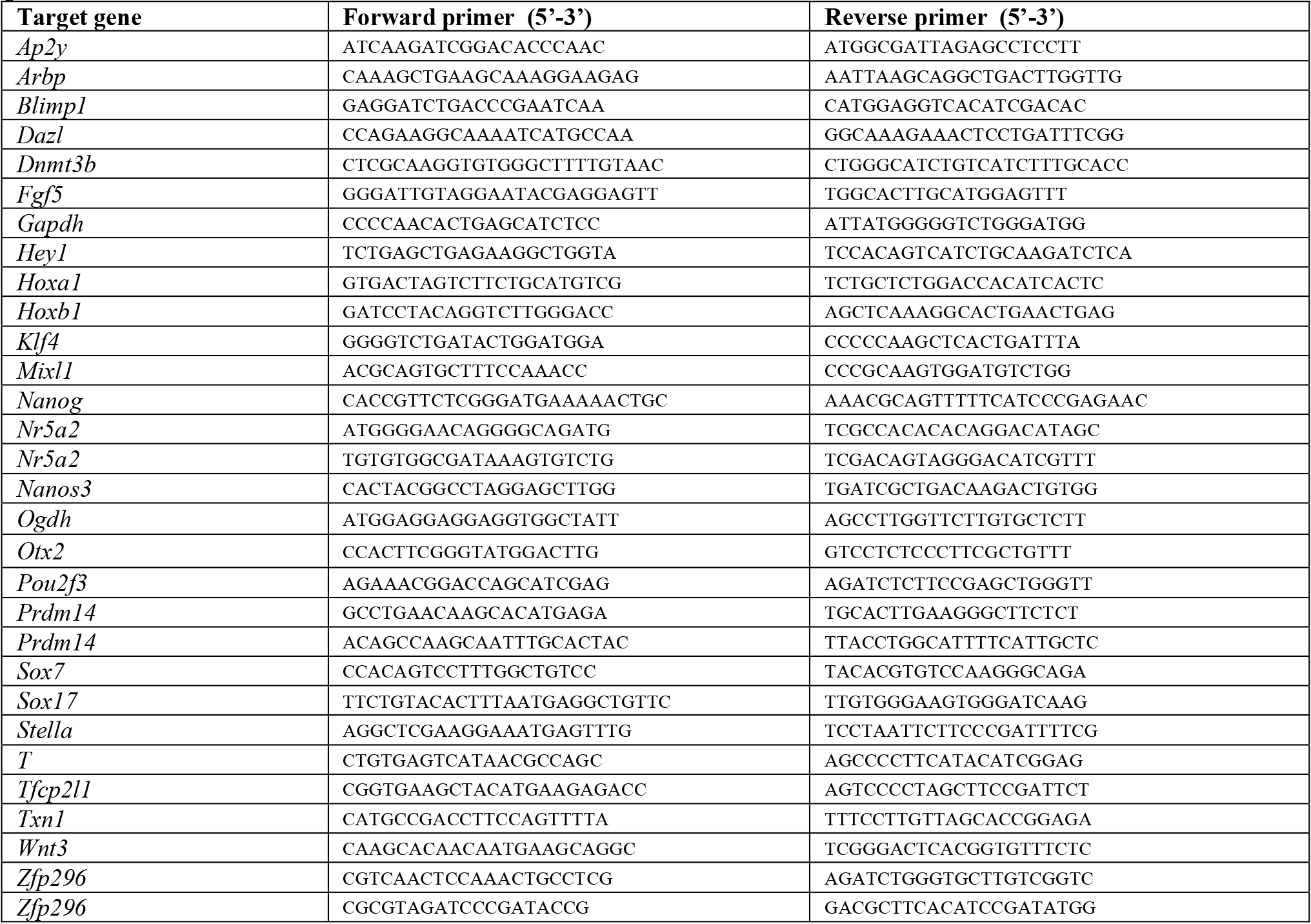

#### gRNA sequences and KO strategy

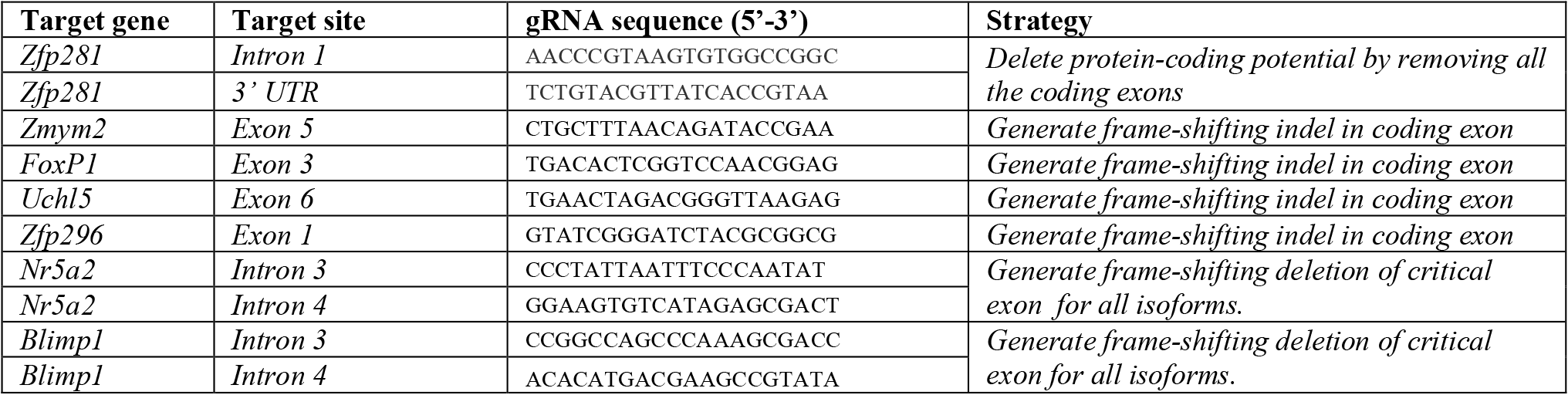

#### CRISPR library amplification including Illumina sequences

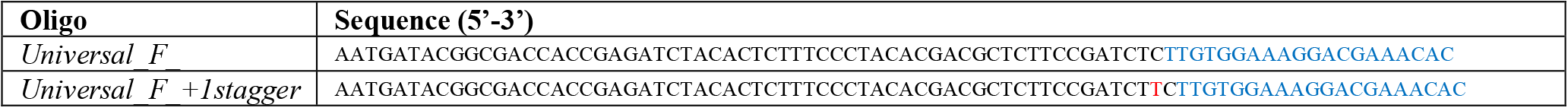

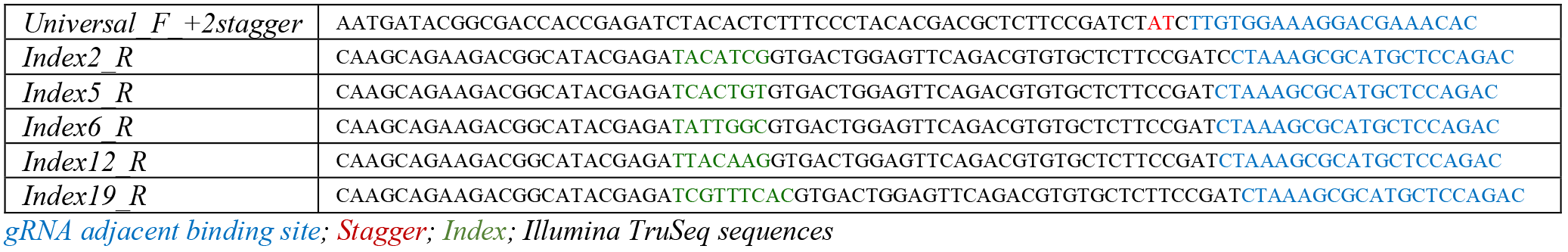

#### siRNA

*gRNA adjacent binding site; Stagger; Index; Illumina TruSeq sequences* *siRNA*
| Target gene | Code | KD efficiency (mRNA) in ESC |
| --- | --- | --- |
| *Ogdh* | ON-TARGETplus siRNA *Ogdh* SmartPool (L-044219-01-0005) | 76% |
| *Txn1* | ON-TARGETplus siRNA *Txn1* SmartPool (L-062799-01-0005) | 84% |

